# Bird species with color polymorphism have greater ecological success

**DOI:** 10.1101/2025.06.19.660624

**Authors:** Kazuya Hata, Eri Sato, Çağan H. Şekercioğlu, Suzuki Noriyuki, Masashi Murakami, Yuma Takahashi

## Abstract

The explanation of the variation in ecological success of species is one of the key challenges in ecology, evolutionary biology and conservation biology. Here, we discuss the function of polymorphism in body color which is one of the most conspicuous features of organisms strongly affecting the performance of species. We applied a phylogenetic comparative approach to assess whether body color polymorphisms contribute to species success in birds, reflected in distributional ranges, extinction risk, and current population trends. We compiled and analyzed a comprehensive global dataset that included the distribution, phylogeny, ecology and morphological traits of all extant bird species. Species with body color polymorphism showed larger distributional ranges, larger environmental ranges, lower risk of extinction, and more stable population trends. Our analyses suggest that the presence of color polymorphism contributes to avian species success at the global scale, especially reflected in lower extinction likelihood and better population trends.

## Introduction

There is considerable variation in ecological success among species (1). The explanation of such variation is one of the challenges in ecology, evolutionary biology and conservation biology (2–5). Species success is often defined using four criteria: 1) more individuals are generated through time, 2) broad niche breadth or adaptive zone, 3) high population density, and 4) large geographic range. Therefore, species success can be evaluated by comparing the variation in range size or extinction risk among species, with the caveat that various ecological attributes should affect species success.

Color polymorphism is expected to be important for the ecological success of a species because different phenotypes are often associated with success at different stages of the life history of a species (3). Several previous studies reported the positive effects of within species genetic or plastic variation on species performance, including productivity, extinction risk or evolutionary potential of the populations (6–9). Accepted explanations for this pattern include niche broadening and the reduction of intraspecific competition (10–13). Body color is one of the best-known examples among ecological traits, which both directly and indirectly affect the species performance (14, 15). For example, the effects of coloration on mate choice (16) and the importance of cryptic coloration for camouflage (17) are well known. Coloration may also have important physical functions, such as the regulation of body temperature and desiccation (18–21). The functional importance of body coloration implies a strong effect of within species color polymorphism on the ecological success, primarily through the modification of interspecific competition or the reduction of predation risk (15).

The positive effects of within species color polymorphism on species distributions have been reported for a variety of organisms, including vertebrates such as amphibians, reptiles and birds, and also invertebrates including butterflies, moths, and damselflies (8, 22– 27). However, these previous studies have lacked a taxonomically comprehensive analysis and have not considered the phylogenetic effects on the species success (28). In addition, the effects of the various life history traits should also be analyzed simultaneously when the effects of color polymorphism on ecological success are tested because various species traits can potentially affect ecological success in general (29). Therefore, to evaluate the role of color polymorphism on species success, it is necessary to focus on taxa that have both comprehensive phylogenetic information and a life history trait database at the global scale.

In the present study, we focused on the world’s bird species to analyze the relationship between color polymorphism and species success. We can utilize variety of information for birds, including distribution (30), phylogeny (31), ecology, and morphological (32) datasets. Furthermore, 3.5% bird species exhibit color polymorphism, without any significant bias in specific taxa (33), which is another advantage for a comprehensive analysis. Here we used a phylogenetic comparative approach to assess whether color polymorphisms contributed to the success of bird species, as reflected in their distributional range and extinction risk.

## Methods

The data set for the presence of color polymorphism in each bird species was derived from the Ducatez et al. (34). Here, the presence of color polymorphism was defined as the occurrence of multiple discrete color patterns within a single population that results directly from genetic variation, thus sexual color polymorphism or geographically clinal color variations were not included in this criterion. This is a very conservative definition of the presence of color polymorphism, thus our evaluation of the effects of color polymorphism on the ecological success of species less likely suffer from type-2 error. In this study, we analyzed these effects on the ecological success of species by concurrently examining the effects of the other factors which might potentially affect the species success, e.g., body size or migratory behavior. Species’ range size, niche breadth, and extinction risk were evaluated as surrogates for the species success.

We used the distributional data of 5,762 terrestrial and freshwater bird species based on their global breeding range maps by Birdlife International (30). The marine bird species were excluded from the analysis because it is difficult to determine the distribution range for these species. Distribution range size (km^2^) of each bird species was calculated on ArcGIS 10.1 (ESRI, Redlands, CA, USA). Niche breadths of the species were evaluated by the zoogeographic regions defined by Holt et al. (35) and also by the environmental variables including meteorological and vegetational ranges. Temperature and precipitation data were taken from the Worldclim2 (36). We also examine the altitudinal ranges available at the USGS site (https://www.usgs.gov/centers/eros/science/usgs-eros-archive-products-overview?qt-science_center_objects=0#qt-science_center_objects). As the measures of vegetation productivity, Normalized Difference Vegetation Index (NDVI) and actual evapotranspiration (AET) were utilized. NDVI data was acquired from ORNL DAAC (37; http://daac.ornl.gov/). We also used the AET map (37) in our analysis. We overlapped the range maps of each bird species with the maps of each variables on ArcGIS. For the zoogeographic regions, the number of regional categories were calculated. For the other environmental variables, maximum and minimum values within their distributional ranges were taken for each bird species. The risk of extinction was evaluated by the IUCN category and the population trends of the species (38). The categories in the IUCN Red List of Threatened Species were converted to a binary index; least concern and near threatened species were converted to 1, and vulnerable, endangered and critically endangered species were converted to 0. The population trends were also converted to a binary index, increasing and stable trends as 1, and decreasing trends as 0. For the Black-faced sheathbill (*Chionis minor*) which distributed in subantarctic islands in the southern Indian Ocean, the data of zoogeographic region, NDVI, and AET were not available, so this species was excluded from the analysis on these variables.

To categorize the bird species, we compiled extensive data on ecologically important trait variables that describe the functional roles of the species (39, 40). We used three trait categories: diet, migratory behavior, and body size. Diet and migratory behaviour of each species were determined from a global bird ecology database that covers all the bird species of the world (32) and is regularly updated with Del Hoyo et al. (41) and other publications. Dietary guilds were classified into seven categories: herbivores including frugivores and nectarivores, granivores, invertebrate eaters, omnivores, and carnivores (including scavenger and piscivores). Smaller body sizes and shorter generation times usually result in faster population growth, allowing populations to maintain higher population densities or larger distributional ranges (42, 43). Migratory behavior was classified as sedentary, latitudinal migration (yearly, regular, long-distance movement) and altitudinal migration (regular movements from high to low altitudes or vice versa based on seasons). Body size of each species was obtained from Dunning (4) and Olson et al. (45) and log-transformed.Phylogenetic logistic regression models were performed with “phyloglm” in phylolm packages in R version 3.5.2 (46). Then we applied a phylogenetic correction to our analyses using phylogenetic generalized linear models (PGLMs), and details can be found in SI Text.

## Results

After accounting for phylogeny, color polymorphism has a positive effect on the distributional range size of a species, the range of zoogeographic regions it is found in, and its environmental niche breadth (Figure 1, SI Appendix, Table S1). No effect of color polymorphism was detected on the range of actual evapotransporation (AET) in a species range. In the best model explaining the range size variation among bird species, migratory status and diet type were both selected (SI Appendix, Table S1). For the models on range of zoogeographic regions, temperature range, and altitudinal range, insectivores had narrower distributional ranges in comparison to the species in other foraging guilds. No effects of activity time were detected in any models (SI Appendix, Table S1).

**Figure 1.**
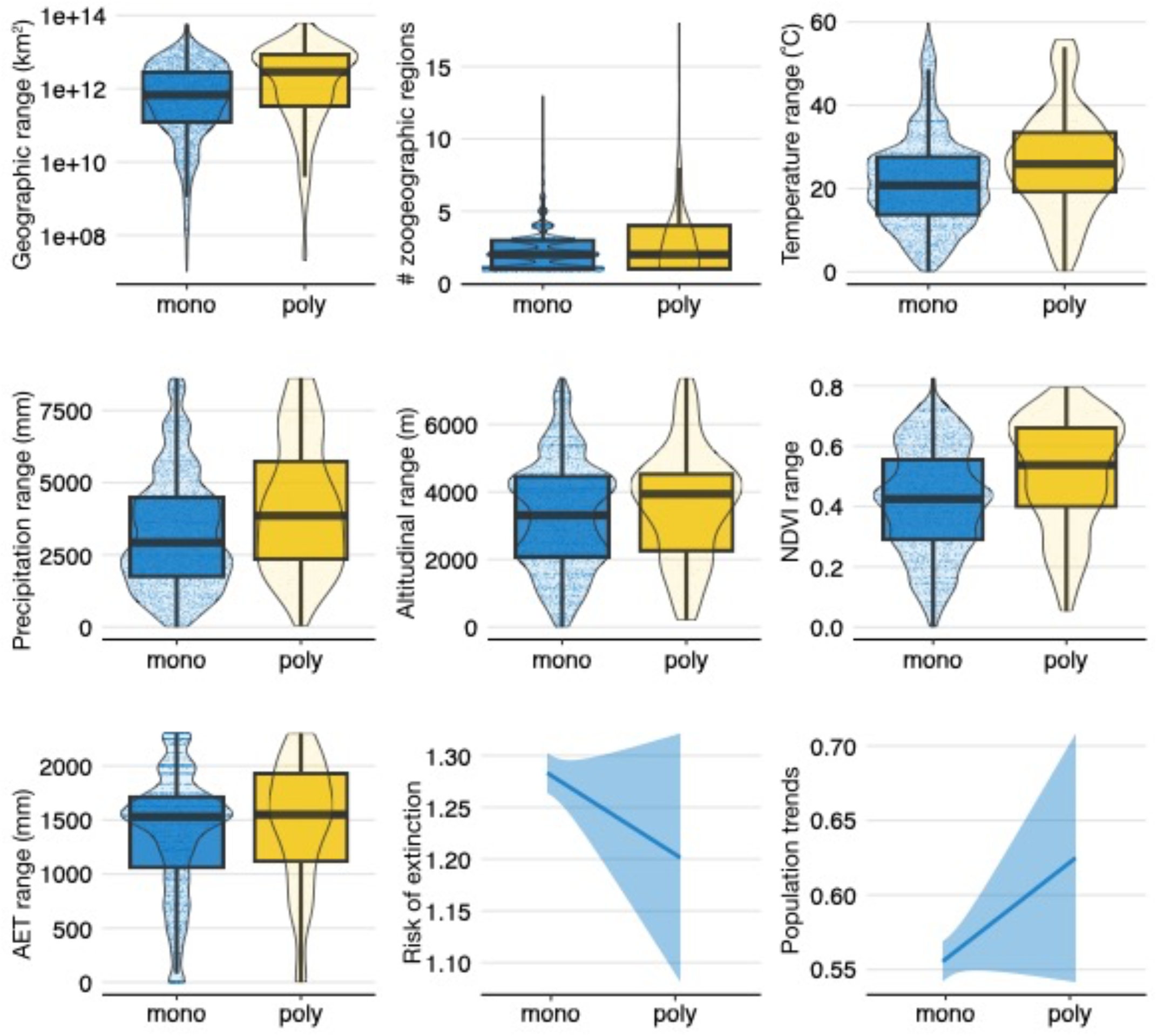
The effect of the color polymorphism on global distribution, extinction risk, and current population trend of bird species. # zoogeographic regions mean the number of regional categories covered by the species.

Species without color polymorphism had higher extinction risk. There was an effect of activity time on extinction risk (SI Appendix, Table S2), but not on the distributional range. Altitudinal migrants had lower extinction risk, whereas latitudinal migrants had higher extinction risk. Color polymorphism also had an effect on population trends (SI Appendix, Table S2), with the species without color polymorphism showing more declines. The effects of body size, migratory trends and diet were included in the best model that explained the variation in extinction risk. The direction of the effects of these explanatory variables on the population trends were mostly identical with those on IUCN categories.

These trends were robust even when the analyses were separately carried out on the species of each bird order. For the five bird orders on which the separate analyses could be carried out (see SI Appendix, Table S3), the general trends, namely positive effects of color polymorphism on the ecological success of species, were robust.

## Discussion

Our results showing the positive effects of color polymorphism on avian species success are global in scale and account for the effects of phylogeny. Furthermore, by simultaneously analyzing the other factors which might potentially affect the ecological success of avian species, we were able to show the contributions of color polymorphism on species success. Species with color polymorphism have broader distributional ranges, lower risk of extinction, and more stable or increasing population trends.

Species characteristics other than color polymorphism had varying effects on species success. For example, body size had a positive effect on a species’ environmental range (SI Appendix, Table S1), probably because larger species can tolerate a broader range of environmental conditions. In contrast, the effect of body size was positive on extinction risk and negative on population trends (decreasing trend in larger species; SI Appendix, Table S1). Larger species are more vulnerable to extinction, have more decreasing population trends (47) and tend to be listed in higher extinction risk categories (48). Among the environmental range variables examined in this study, AET was the only variable on which no effect of color polymorphism was detected. Because AET is a function of both environmental and vegetation factors such as temperature, precipitation and NDVI (49), on which we observed positive effects of color polymorphism in our analyses, the reason for this lack of effects is difficult to interpret.

The direction of causality between color polymorphism and large-scale ecological patterns such as distributional ranges is not straightforward to examine. Species with greater distributions are likely to be exposed to more variable environmental conditions and may therefore have more intraspecific phenotypic variation than those with narrow distributions (50). In fact, one of the common mechanisms for the maintenance of genetic polymorphism within a population is considered to be the combination of spatially varying selection and gene flow through dispersal (51–55). It suggests that larger distributions and environmental ranges result in color polymorphism within a population, not the other way around. However, in our study, we examined the effect of color polymorphism on extinction risk by simultaneously considering geographical range as a covariate. There was still a significant relationship between color polymorphism and lower extinction risk (SI Appendix, Table S2), indicating increased species success. Therefore, our results suggest that extinction risk may be mitigated through the positive effects of color polymorphism, even after accounting for geographical range size. In addition, population trend is another indicator of species success that is independent from range size and is observed at ecological timescales of ∼50 years (53). For birds, which have relatively long generation times, it is reasonable to assume that current population trend is unlikely to affect the evolution of color polymorphism at such a short timescale, suggesting that it is the presence of color polymorphism that has an impact on current population trend.

Phenotypic variation within species has a range of consequences for the evolution and ecological performance of populations and species (56). For example, Takahashi et al. (13) showed the positive diversity effect of genetic polymorphism on the population productivity attributed by the complementarity effect in *Drosophila* flies. Sultan (57) showed that developmental plasticity in plant functional traits decreases extinction risk of species.Summarizing these results, Wennersten and Forsman (58) highlighted the importance of spatial and temporal scales to elucidate the mechanisms and the consequences of phenotypic variation on species success. For a more detailed examination of the link between species success and within species polymorphism, further investigations of the consequences of polymorphism should facilitate the understanding of the ecological dynamics of natural populations and communities.

Our findings on the ecological consequences of color polymorphism can also be applied to the conservation management of wild birds and other organisms. Various studies on extinction risk of species have focused on testing the relative importance of intrinsic biological factors to predict extinction risk, and others have shown that anthropogenic threats are equally important (59, 60). Our results show the importance of color polymorphism of species as another predictor of extinction risk, indicating the increased vulnerability of monomorphic species faced with global changes. In addition, Forsman (61) showed the higher probability of colonization success for polymorphic species compared with monomorphic species. This further reduces the extinction likelihood of polymorphic species and also suggests that polymorphic species have a higher risk of becoming invasive.

## ACKNOWLEDGEMENTS

We thank the member of Functional Ecology Lab in Chiba University for their assistance. Funding was provided by the JSPS (to T.Y., M.M., and E.S.) and the Toyota Foundation and Sumitomo Foundation (to Y.T). Ç.H.Ş thanks H. Batubay Özkan and Barbara Watkins for their support of the Biodiversity and Conservation Ecology Lab at the University of Utah School of Biological Sciences. We are grateful to dozens of volunteers and students, especially Monte Neate‐Clegg, Joshua Horns, Evan Buechley, Jason Socci, David Blount, Sherron Bullens, Debbie Fisher, David Hayes, Beth Karpas and Kathleen McMullen, for their dedicated help with BirdBase, the world bird ecological trait database.

