## Appendix for "Bird species with color polymorphism have greater ecological success"

This PDF file includes:
- Supplementary text (SI Methods)
- Tables S1 to S7
- Legends for Datasets S1
- SI References

Other supplementary materials for this manuscript include the following:
- Datasets S1

**Supplementary Information Text**

### SI Methods

Phylogenetic logistic regression models were performed with "phyloglm" in phylolm packages in R version 3.5.2 (1). We used the phylogeny of extant birds (2). Then we applied a phylogenetic correction to our analyses using phylogenetic generalized linear models (PGLMs). The effect of the color polymorphism on the ecological success of species was evaluated by model selection. Full models were included the "color polymorphism", "body size", "longevity", "migratory behavior" and "diet type" as the explanatory variables. For the analysis on the extinction risks, we also included the geographical range as a covariate. We used the backward stepwise deletion of the variables from the full model based on the AIC.

First, we applied this analysis on the 5,764 terrestrial and freshwater bird species (Table S1, S2). Then, we carried out separate analyses on the species of each bird orders. Following the number of bird species with color Polymorphism in each bird order, we selected five bird orders for the analyses (Table S3, S4, S5).

### SI Tables

Table S1. The effects of the color polymorphism on the ecological success of species were analyzed with examining the other factors potentially affects the success. The ecological success of species was evaluated by species range size and niche breadth. In these analyses, the values between first and third quantiles were utilized to evaluate the effects of outliers in each species range. The coefficients of each explanatory variable selected in best model based on AIC. The numbers in parentheses show the number of models where this trait was included in the best model among 100 phylogenetic trees based on AIC. ΔAIC 1 showed ($AIC_{null} - AIC_{best}$). ΔAIC 2 shows the difference between the AIC value of the best model without polymorphism and that of the best model ($AIC_{best\_w/o\_poly} - AIC_{best}$).

| traits | | Temperature (°C) | Precipitation (mm) | Elevation (m) | NDVI | AET (mm) |
|---|---|---|---|---|---|---|
| color polymorphism | | 0.40 – 0.49 (100) | – (***100) | – (***100) | 0.010 – 0.012 (100) | 44.1 – 51.7 (27) |
| body size | | 0.10 – 0.14 (100) | – | 14.6 – 18.4 (56) | 0.0019 – 0.0022 (17) | −23.4 – −14.4 (56) |
| longevity | | – | −14.8 – −4.0 (100) | – | 0.00097 – 0.00103 (4) | −11.8 – −7.3 (100) |
| migration* | altitudinal | 2.4 – 2.5 (100) | 60.6 – 67.6 (100) | 461 – 485 (100) | 0.038 – 0.039 (100) | 18.0 – 34.2 (100) |
| | latitudinal | 2.4 – 2.5 | −96.3 – −90.6 | 52.3 – 755 | 0.044 – 0.045 | −118 – −105 |
| diet** | frugivore | −0.11 – −0.18 (100) | 54.0 – 64.3 (100) | −10.9 – 7.4 (100) | −0.0093 – −0.0072 (100) | 35.4 – 50.1 (100) |
| | omnivore | 0.15 – 0.21 | 2.55 – 9.56 | 21.8 – 36.9 | 0.0018 – 0.0035 | −4.61 – 8.19 |
| | granivore | 0.72 – 0.84 | −73.1 – −58.0 | 73.4 – 96.4 | 0.022 – 0.024 | −62.4 – −33.4 |
| | carnivore | 1.2 – 1.7 | −6.3 – 4.2 | 23.2 – 53.6 | 0. 027– 0.031 | −52.0 – −23.2 |
| ΔAIC_1 | | 745 – 793 | 71.7 – 85.0 | 214 – 240 | 391 – 407 | 63.4 – 81.4 |
| ΔAIC_2 | | 1.8 – 2.6 | – | – | 1.7 – 2.6 | −0.03 – 0.52 |

* sedentary species were set as the reference (= 0)
** insectivores were set as the reference (= 0)
*** the number of models where this trait was included in the models with ΔAIC < 2 of best models among 100 phylogenetic trees

Table S2. Number of species with/without color polymorphism for each of 5 dominant bird orders and the other orders.

| bird order | non Polymorphism | Polymorphism |
|---|---|---|
| Accipitriformes | 135 | 30 |
| Caprimulgiformes | 341 | 16 |
| Cuculiformes | 73 | 12 |
| Passeriformes | 3811 | 40 |
| Strigiformes | 61 | 25 |
| the other orders | 1194 | 26 |

Table S3. The effects of the color polymorphism on the ecological success of species were analyzed for 5 bird orders with examining the other factors potentially affects the success. The ecological success of species was evaluated by species range size and niche breadth. The range of the coefficients for the color polymorphism in best model based on AIC were shown with the number in parentheses of show the number of models where color polymorphism was included in the best model among 100 phylogenetic trees.

| bird order | Geographic Range ($km^2$) | Zoogeographic Regions | Temperature (°C) | Precipitation (mm) | Elevation (m) | NDVI | AET (mm) |
|---|---|---|---|---|---|---|---|
| Accipitriformes | – | – | – | – | 419.5 – 420.8 (100) | 0.064 − 0.068 (100) | – |
| Caprimulgiformes | – | – | – | – | −684.7 – − 580.1 (100) | −0.060 – −0.049 (80) | 182.3 – 226.3 (97) |
| Cuculiformes | – | – | – | 1195.0 – 1293.6 (100) | – | – | – |
| Passeriformes | 0.87 – 0.98 (100) | 0.87 – 0.99 (100) | 4.00 – 4.57 (100) | 843.4 – 949.1 (100) | 334.9 – 408.1 (100) | 0.071 − 0.076 (100) | 126.2 − 154.1 (100) |
| Strigiformes | 0.94 – 1.15 (29) | 0.93 – 1.15 (31) | – | – | – | 0.044 − 0.071 (5) | – |

Table S4. The effects of the color polymorphism on the ecological success of species were analyzed for 5 bird orders with examining the other factors potentially affects the success. The ecological success of species was evaluated by extinction risks. The range of the coefficients for the color polymorphism in best model based on AIC were shown with the number in parentheses of show the number of models where color polymorphism was included in the best model among 100 phylogenetic trees.

| traits | risk of extinction | population trend |
|---|---|---|
| Accipitriformes | 1.63 – 1.69 (19) | 0.25 – 0.90 (28) |
| Caprimulgiformes | – | −0.16 (1) |
| Cuculiformes | – | – |
| Passeriformes | 0.63 – 0.68 (32) | 0.87 – 0.99 (25) |
| Strigiformes | – | 0.93 – 1.15 (3) |

**Dataset S1 (separate file).** Bird trait data used in this study.
